# Regulation of bacterial cell cycle progression by redundant phosphatases

**DOI:** 10.1101/2020.04.29.069633

**Authors:** Jérôme Coppine, Andreas Kaczmarczyk, Kenny Petit, Thomas Brochier, Urs Jenal, Régis Hallez

## Abstract

In the model organism *Caulobacter crescentus*, a network of two-component systems involving the response regulators CtrA, DivK and PleD coordinate cell cycle progression with differentiation. Active phosphorylated CtrA prevents chromosome replication in G1 cells while simultaneously regulating expression of genes required for morphogenesis and development. At the G1-S transition, phosphorylated DivK (DivK~P) and PleD (PleD~P) accumulate to indirectly inactivate CtrA, which triggers DNA replication initiation and concomitant cellular differentiation. The phosphatase PleC plays a pivotal role in this developmental program by keeping DivK and PleD phosphorylation levels low during G1, thereby preventing premature CtrA inactivation. Here, we describe CckN as a second phosphatase akin to PleC that dephosphorylates DivK~P and PleD~P in G1 cells. However, in contrast to PleC, we do not detect kinase activity with CckN. The effects of CckN inactivation are largely masked when PleC is present, but become evident when PleC and DivJ, the major kinase for DivK and PleD, are absent. Accordingly, mild overexpression of *cckN* restores most phenotypic defects of a *pleC* null mutant. We also show that CckN and PleC are proteolytically degraded in a ClpXP-dependent way well before the onset of the S phase. Surprisingly, known ClpX adaptors are dispensable for PleC and CckN proteolysis, suggesting the existence of adaptors specifically involved in proteolytic removal of cell cycle regulators. Since *cckN* expression is induced in stationary phase, depending on the stress alarmone (p)ppGpp, we propose that CckN acts as an auxiliary factor responding to environmental stimuli to modulate CtrA activity under suboptimal conditions.

**Importance:** Two-component signal transduction systems are widely used by bacteria to sense environmental signals and respond accordingly by modulating various cellular processes, such as cell cycle progression. In *Caulobacter crescentus*, PleC acts as a phosphatase that indirectly protects the response regulator CtrA from premature inactivation during the G1 phase of the cell cycle. Here, we provide genetic and biochemical evidence that PleC is seconded by another phosphatase, CckN. The activity of PleC and CckN phosphatases is restricted to G1 phase since both proteins are timely degraded by proteolysis just before the G1-S transition. This degradation requires new proteolytic adaptors as well as an unsuspected N-terminal motif for CckN. Our work illustrates a typical example of redundant functions between two-component proteins.

## Introduction

The α-proteobacterium *Caulobacter crescentus* divides asymmetrically to generate two daughter cells with different cell fates, a sessile stalked cell and a motile swarmer cell. While the newborn stalked cell can immediately re-enter S phase and initiate chromosome replication, the smaller swarmer cell engages in an obligatory motile and chemotactic but non-replicative G1 phase. Concomitantly with its entry into the S phase (G1-S transition), the swarmer cell differentiates into a stalked cell (swarmer-to-stalked cell transition). A complex regulatory network controlling the activity of the central and essential response regulator CtrA coordinates different cell cycle stages with accompanying morphological changes and development. CtrA activity is carefully regulated throughout the cell cycle at the transcriptional and post-translational levels. CtrA protein levels and its phosphorylation status are mostly determined by the action of a phosphorelay involving the hybrid kinase CckA and its cognate histidine phosphotransferase ChpT (1–4). In the swarmer cell, the kinase activity of CckA is stimulated at the flagellated pole by the physical contact with the non-conventional histidine kinase DivL (5–8). DivL is free to activate CckA since its inhibitor – the response regulator DivK – is dephosphorylated (*i.e.* inactivated) by the phosphatase PleC (PleC^P^). Hence, CckA promotes the ChpT-dependent phosphorylation of CtrA, thereby stimulating its activity. At the same time, the CckA/ChpT phosphorelay also protects CtrA from its proteolytic degradation by phosphorylating CpdR, a response regulator whose the unphosphorylated form primes the ClpXP protease for CtrA degradation (4, 9). Active CtrA (CtrA~P) binds the single chromosomal origin of replication (*C*_*ori*_) to prevent DNA replication initiation (Figure 1a). As a transcription factor, CtrA~P also directly activates or represses the expression of more than 200 genes involved in multiple biological processes including cell cycle, cell differentiation and cell division (10).

**Figure 1:**
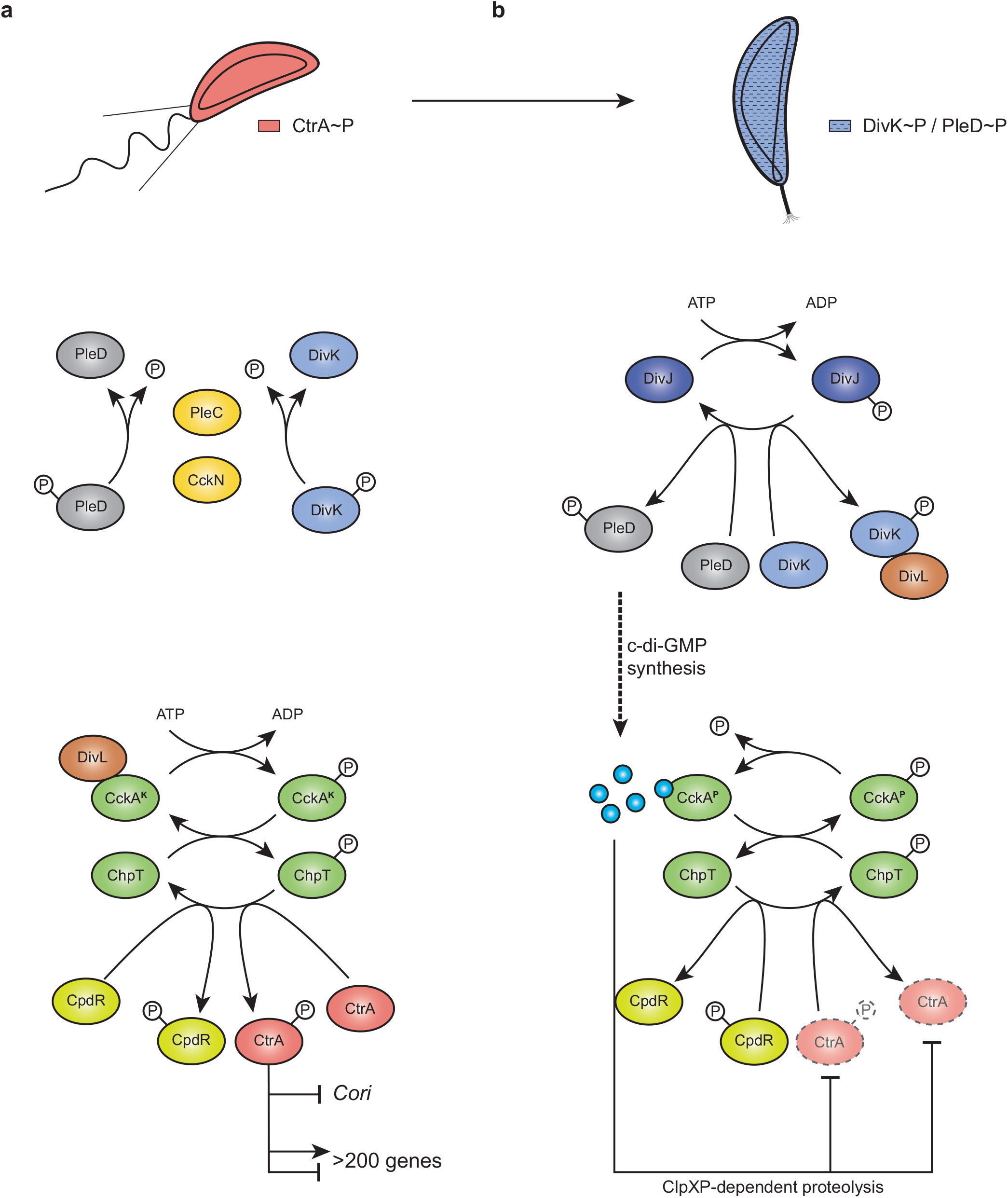
CtrA regulation pathway in *Caulobacter crescentus* in (a) swarmer and (b) stalked cells. In swarmer cells (a), DivK is actively dephosphorylated by PleC and CckN, and therefore not able to interact with DivL. Free DivL activates the phosphorelay culminating in CtrA and CpdR phosphorylation. Active CtrA (CtrA~P) regulates the expression of more than 200 genes and inhibits DNA replication initiation by binding the single chromosomal origin of replication (*Cori*). At the G1-S transition (b), CckN and PleC are cleared form the cells while DivK and PleD are phosphorylated by their cognate histidine kinase DivJ. Phosphorylated DivK (DivK~P) interacts with DivL and reduces its affinity for CckA leading to an inhibition of its kinase activity on CtrA and CpdR. Phosphorylation of PleD promotes its diguanylate cyclase activity resulting in an increased synthesis of c-di-GMP. High levels of c-di-GMP not only stimulates CckA phosphatase activity on both CpdR~P and CtrA~P, but also drives, concomitantly with unphosphorylated CpdR, ClpXP-dependent degradation of CtrA. Together, this results in the rapid inactivation of CtrA allowing DNA replication initiation to proceed.

At the G1-to-S transition, DivK becomes highly phosphorylated. This results from (i) a switch from PleC phosphatase to kinase activity before proteolytic removal of PleC and (ii) post-translational stimulation of DivJ, the major histidine kinase responsible for DivK and PleD phosphorylation (11, 12). Once phosphorylated, DivK~P physically interacts with DivL and strongly reduces its affinity for CckA (7, 8, 13). Hence, the kinase activity of CckA is no longer stimulated. Simultaneously, CckA phosphatase activity is directly stimulated by c-di-GMP, the levels of which strongly and rapidly raise due to activation of the diguanylate cyclase PleD and inactivation of the phosphodiesterase PdeA. PleD becomes highly phosphorylated (*i.e.* activated) by DivJ at the differentiating pole (14, 15), whereas PdeA is degraded by ClpXP (16). High levels of c-di-GMP also drive ClpXP-dependent degradation of CtrA directly by binding to the proteolytic adaptor PopA (17, 18). Together, these events result in the rapid inactivation of CtrA during G1-S transition and trigger an irreversible program leading to chromosome replication and cell differentiation (Figure 1b). Inactivation of PleC phosphatase activity and the resulting increase in DivK~P (and PleD~P) are the earliest known events in this G1-S transition signaling pathway. We thus wondered whether other factors besides PleC and DivJ could influence DivK and PleD (de)phosphorylation.

Almost 20 years ago, interaction partners of DivK was identified in a yeast two-hybrid screen (13). Apart from PleC, DivJ and DivL, which were unsurprisingly found as prominent hits, another histidine kinase called CckN was discovered in this study as a physical partner of DivK (13), but the role played by this actor in the CtrA regulatory network has not been characterized so far. Here we show that similarly to PleC, CckN displays phosphatase activity towards DivK and PleD during the G1/swarmer phase of the cell cycle. However, in contrast to PleC, the kinase activity of CckN cannot be activated by DivK~P at the G1-S transition. Both phosphatases are required to sustain optimal CtrA activity in the non-replicative swarmer cells before being inactivated by proteolysis at the G1-S transition. Interestingly, we also show that both CckN and PleC are the earliest CtrA regulatory proteins to disappear, likely by ClpXP-dependent proteolysis, suggesting that the concomitant degradation of both phosphatases might be the signal initiating cell cycle entry. Surprisingly, these degradations do not rely on any known proteolytic adaptors. In addition, we show that *cckN* expression is stimulated in stationary phase, depending on (p)ppGpp. We propose a model in which CckN influences CtrA activity under non-optimal growth conditions.

## Results

### CckN is a second phosphatase for DivK and PleD

CckN was previously identified as an interaction partner of DivK in a yeast two-hybrid screen (13). The interaction of CckN with DivK was confirmed by coimmunoprecipitation (Figure 2a) and bacterial two-hybrid assays (Figure 2b). We next tested whether CckN displayed kinase activity, *i.e.* could autophosphorylate *in vitro* in the presence of ATP. Purified CckN with either a N-terminal His6 or a His6-MBP tag did not show autokinase activity *in vitro* in our experiments, despite the presence of a predicted HATPase domain (pfam02518) and all the catalytic residues in the DHp and CA domains conserved in prototypical HisKA-type histidine kinases (19). In contrast, we detected robust autophosphorylation of His6-MBP-tagged purified *C. crescentus* DivJ and PleC proteins encompassing the cytoplasmic catalytic histidine kinase core region as well as of His6-tagged purified DivJ comprising the soluble cytoplasmic catalytic core region from *Sinorhizobium meliloti* (DivJ^Sm^) (Figure 2c, Supplementary Figure 1a). A non-phosphorylatable variant of DivK (DivK_D53N_) stimulated *C. crescentus* DivJ and PleC autokinase activity, as reported before (15), but did not show any stimulatory effect on CckN autophosphorylation (Supplementary Figure 1a). Accordingly, DivK could be phosphorylated with the non-cognate DivJ^Sm^, but not with CckN. In contrast, CckN was able to efficiently dephosphorylate DivK~P (Figure 2c). Since PleC and DivJ are known to also (de)phosphorylate another response regulator akin to DivK, PleD, we next tested whether CckN could dephosphorylate PleD. As shown in Figure 2d, CckN was able to rapidly dephosphorylate PleD. In the presence of DivK_D53N_ in reactions containing DivJ and CckN, PleD dephosphorylation was still observed (Supplementary Figure 1b), suggesting that in contrast to PleC, the kinase/phosphatase balance of CckN is not regulated by DivK_D53N_. Finally, we measured PleD~P levels *in vivo* in strains overexpressing *cckN*. In line with *in vitro* data, we found that the phosphorylated form of PleD (PleD~P) was strongly reduced upon overexpression of wild-type *cckN* compared to a control strain, whereas overexpression of a mutant variant of *cckN* (*cckN*_*H47A*_) that has the predicted phosphorylatable His47 residue mutated to Ala did not influence PleD~P levels (Figure 2e). Together, these results suggest that CckN is a phosphatase but not a kinase for both DivK and PleD.

**Figure 2:**
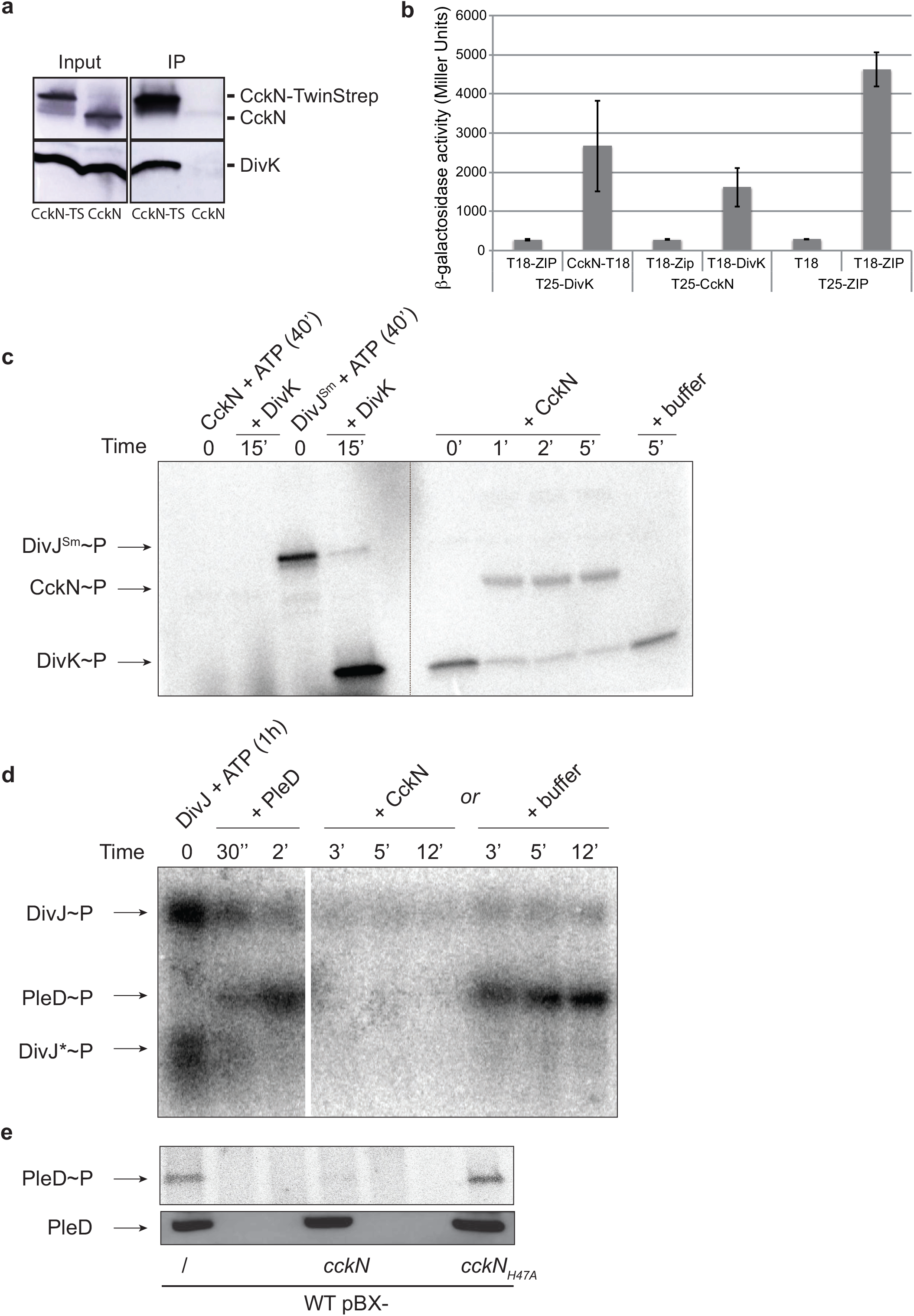
CckN is a phosphatase for DivK and PleD. (a) Co-immunoprecipitation (Co-IP) experiments showing that CckN and DivK are part of the same protein complex. Co-IP were performed on protein extracts of *cckN-TwinStrep* (RH2235) and wild-type (RH2070) strains. CckN and DivK were detected by Western blotting using respectively anti-CckN and anti-DivK antibodies before (Input) and after immunoprecipitation (IP) with Strep-Tactin coated magnetic beads. CckN-TS: CckN-TwinStrep (b) Bacterial two-hybrid assay showing that CckN and DivK interact with each other. β◻-galactosidase assays were performed on MG1655 *cyaA∷frt* (RH785) strains coexpressing T18 fused to *ZIP*, *cckN*, or *divK* with T25 fused to *ZIP*, *cckN*, or *divK*. T18 and T25 alone were used as negative controls while coexpression of T18-ZIP and T25-ZIP was used as a positive control. Error bars = SD, n ≥ 3. (c) *In vitro* phosphorylation assay showing that CckN cannot phosphorylate DivK but can dephosphorylate DivK~P. CckN or DivJ^Sm^ was incubated alone for 40’ with [γ-^32^P]ATP before adding DivK for 15’. Then, DivK phosphorylated by DivJ^Sm^ was washed to remove excess of [γ-^32^P]ATP (dotted line) and incubated with or without CckN for the indicated time. (d) *In vitro* phosphorylation assay showing that CckN can dephosphorylate PleD~P. DivJ was incubated alone for 1h with [γ-^32^P]ATP before adding PleD for 2’ before adding CckN or buffer incubating for the indicated time. DivJ* likely corresponds to a degradation product of DivJ (e) *In vivo* phosphorylation assay showing that overexpression of functional *cckN* decreases of PleD phosphorylation. Wild-type (RH50) cells harbouring the pBX, pBX-*cckN* or *cckN*_*H47A*_ plasmid were grown for 3 h in M5G with 0.05 mM phosphate supplemented with 0.1% xylose. The phosphorylation (up) and protein (down) levels of PleD were determined *in vivo* as described in the Material & Methods.

### CckN impacts CtrA activity through DivK

As a regulator of DivK phosphorylation, CckN is expected to affect CtrA activity. To test this hypothesis, we monitored the activity of CtrA-dependent promoters in different genetic backgrounds using *lacZ* transcriptional reporters and β-galactosidase assays. We found that inactivating *cckN* either in an otherwise wild-type did not change substantially CtrA activity (Supplementary Figure 2a). Likewise, Δ*cckN* did not interfere the hyper-activity of CtrA measured in a Δ*divJ* background (Supplementary Figure 2a). However, the inactivation of *cckN* in a Δ*divJ* Δ*pleC* background strongly diminished activity of CtrA-dependent promoters, but did not influence activity of promoters that are not regulated by CtrA (Figure 3a, Supplementary Figure 2b). The reason why the effects of *cckN* inactivation could only be detected when both DivJ and PleC were absent is likely because PleC constitutes the main phosphatase of DivK and thus needs to be first inactivated to unmask the effects of CckN. However, the activity of CtrA-dependent promoters is so low in Δ*pleC* cells (Supplementary Figure 2a) that *divJ* had to be concomitantly inactivated with *pleC* to study the impact of *cckN* inactivation on those promoters. In support of this notion, we found that Δ*cckN* modulated activity of CtrA in a *pleC*_*F778L*_ background (Supplementary Figure 3a). This PleC variant was described to lack kinase but not phosphatase activity (PleC^K-P+^) and was shown to complement motility, phage sensitivity and stalk biogenesis defects of a Δ*pleC* mutant when expressed on a multicopy plasmid (20). However, we found that CtrA-dependent promoters were less active in *pleC*_*F778L*_ cells – in which *pleC*_*F778L*_ was expressed as the only copy from the endogenous *pleC* locus – than in wild-type cells, but more active than in Δ*pleC* cells (Supplementary Figure 2a, Supplementary Figure 3a). These data suggest that the phosphatase activity of PleC_F778L_ is slightly reduced compared to wild-type PleC. Accordingly, the activity of CtrA-dependent promoters was further decreased in *pleC*_*F778L*_ Δ*cckN* cells to levels observed with fully inactive alleles of *pleC*, *e.g.* Δ*pleC* or *pleC*_*H610A*_ (Supplementary Figure 3a). In line with these effects, *cckN* inactivation in a *pleC*_*F778L*_ background further decreased motility and attachment compared to the parental *pleC*_*F778L*_ strain (Supplementary Figure 3b). Together, these data support the idea that PleC masks the phosphatase activity of CckN on DivK under standard laboratory conditions.

**Figure 3:**
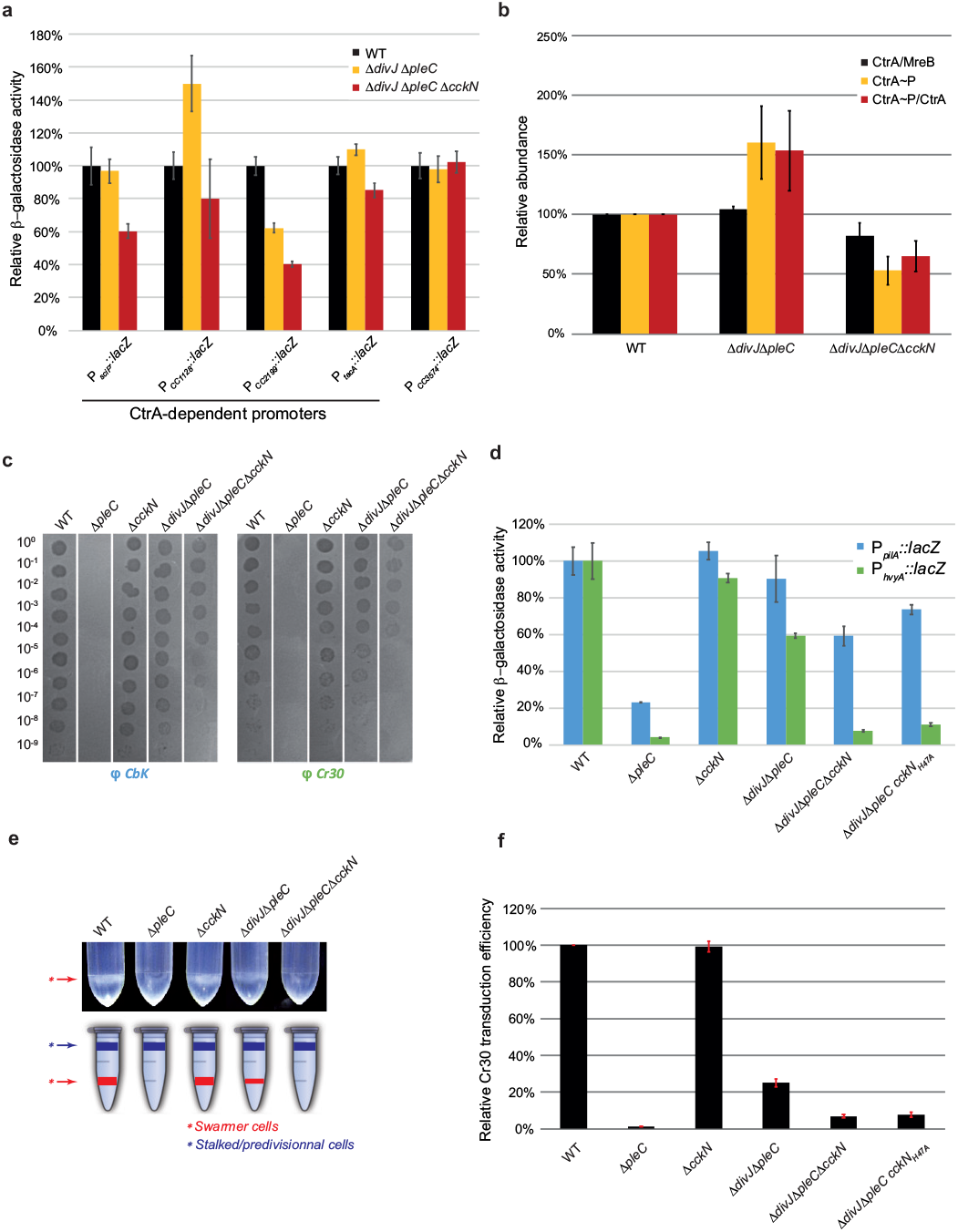
CckN controls development by regulating CtrA activity. (a) β◻-galactosidase assays were performed on wild-type (RH50), Δ*divJ* Δ*pleC* (RH1103) and Δ*divJ* Δ*pleC* Δ*cckN* (RH1111) strains harbouring *lacZ* fusions to CtrA-dependent (P_*sciP*_, P_*CC1128*_, P_*CC2199*_ and P_*tacA*_) and CtrA independent (P_*CC3574*_) promoters, grown in complex medium (PYE) and normalized to the WT (100%). Error bars = SD, n ≥ 3. (b) The protein and the phosphorylation levels of CtrA were measured in wild-type (RH50), Δ*divJ* Δ*pleC* (RH1103) and Δ*divJ* Δ*pleC* Δ*cckN* (RH1111) strains and normalized to the WT (100%). The CtrA protein levels normalized to the MreB levels (black bars) were determined by western blotting. The CtrA phosphorylation levels (yellow bars) were determined *in vivo* as described in the Material & Methods. The CtrA~P/CtrA ratio (red bars) were obtained by dividing black values by yellow values. Error bars = SD, n = 3. (c) Bacteriophages sensitivity assays were performed with CbK and Cr30 on wild-type (RH50), Δ*pleC* (RH217), Δ*cckN* (RH1106), Δ*divJ* Δ*pleC* (RH1103) and Δ*divJ* Δ*pleC* Δ*cckN* (RH1111) strains on PYE agar plates. (d) β◻-galactosidase assays were performed on wild-type (RH50), Δ*pleC* (RH217), Δ*cckN* (RH1106), Δ*divJ* Δ*pleC* (RH1103), Δ*divJ* Δ*pleC* Δ*cckN* (RH1111) and Δ*divJ* Δ*pleC cckN*_*H47A*_ (RH1800) strains harbouring P_*pilA*_*∷lacZ* or P_*hvyA*_*∷lacZ* fusions, grown in complex medium (PYE) and normalized to the WT (100%). Error bars = SD, n ≥ 3. (e) Buoyancy was evaluated by mixing 550 μl of Ludox LS Colloidal Silica (30%) to 1 ml of wild-type (RH50), Δ*pleC* (RH217), Δ*cckN* (RH1106), Δ*divJ* Δ*pleC* (RH1103) and Δ*divJ* Δ*pleC* Δ*cckN* (RH1111) strains grown in PYE. Then, the mix was centrifuged for 15 min at 9,000 rpm. (f) Cr30-dependent transduction efficiency was measured by transducing Cr30 lysates (LHR73 or LHR75) in wild-type (RH50), Δ*pleC* (RH217), Δ*cckN* (RH1106), Δ*divJ* Δ*pleC* (RH1103), Δ*divJ* Δ*pleC* Δ*cckN* (RH1111) and Δ*divJ* Δ*pleC cckN*_*H47A*_ (RH1800) strains grown in complex medium (PYE) and normalized to the WT (100%). Error bars = SD, n = 2.

Given that DivK~P negatively affects CtrA phosphorylation and protein levels (4, 21), we monitored phosphorylation and protein abundance of CtrA in Δ*divJ* Δ*pleC vs* Δ*divJ* Δ*pleC* Δ*cckN* cells. Whereas CtrA abundance did not vary substantially, the phosphorylation state of CtrA was strongly diminished in the triple mutant (Δ*divJ* Δ*pleC* Δ*cckN*) in comparison to the parental Δ*divJ* Δ*pleC* strain (Figure 3b). Thus, CckN impacts CtrA activity mainly by promoting its phosphorylation, presumably via dephosphorylation of DivK. Comparative chromatin immunoprecipitation experiments coupled to deep sequencing (ChIP-seq) in Δ*divJ* Δ*pleC* Δ*cckN* vs Δ*divJ* Δ*pleC* cells showed that *cckN* inactivation impacts the entire CtrA regulon (Supplementary Figure 3c, Supplementary Table 1).

Finally, we checked whether the effect of CckN on CtrA phosphorylation and activity was fully dependent on DivK by monitoring the activity of CtrA in the absence of *divK*. As *divK* is essential but was shown to be dispensable in a *cpdR*_*D51A*_ background (22), we measured the activity of CtrA-regulated promoters in *cpdR*_*D51A*_ Δ*divK* with or without *cckN*. In contrast to *divK*^*+*^ cells (Figure 3a, d), inactivating *cckN* did not influence anymore CtrA activity in the absence of DivK (Supplementary Figure 3d), suggesting that CckN acts on CtrA mostly – if not entirely – through DivK. Altogether, these results suggest that CckN is a phosphatase of DivK akin to PleC, ultimately keeping CtrA active in G1 cells.

### CckN controls development in a CtrA-dependent way

Next, we tested whether *cckN* inactivation displayed developmental defects. A Δ*pleC* strain is known to be fully resistant to both CbK and Cr30 bacteriophages (Figure 3c) (23, 24). This is due to *pilA* and *hvyA* transcription being directly activated by CtrA~P, and not being sufficiently expressed in the absence of PleC (Figure 3d) (24, 25). *pilA* codes for the major pilin subunit of polar pili used by phage CbK as a receptor (26). *hvyA* encodes a transglutaminase homolog that specifically protects swarmer cells from capsulation, thereby allowing the Cr30 bacteriophage to reach its receptor, the S-layer (27). Thus, neither CbK nor Cr30 can infect Δ*pleC* cells since the polar pili are absent and the S-layer in inaccessible. Because the differential capsulation of *Caulobacter* daughter cells is also responsible for their specific buoyancy properties, with the non-capsulated swarmer cells being heavy and the other capsulated cell types light (24), Δ*pleC* swarmer cells lack their typical low buoyancy feature by becoming capsulated (Figure 3e). As expected and as reported before, full resistance to both bacteriophages, high buoyancy and low P_*pilA*_ and P_*hvyA*_ activity displayed by the single Δ*pleC* strain was restored by inactivating *divJ* (Figure 3c-e) (11, 24, 28). Interestingly, we found that a non-functional allele of *cckN* (Δ*cckN*) reduced sensitivity to bacteriophages and lost low buoyancy in a Δ*divJ* Δ*pleC* or *pleC*_*F778L*_ background, but not in an otherwise wild-type background (Figure 3c-e, Supplementary Figure 3e). In line with the lower sensitivity to Cr30 infection, the relative Cr30-mediated transduction efficiency was also reduced in Δ*divJ* Δ*pleC* Δ*cckN* and Δ*divJ* Δ*pleC cckN*_*H47A*_ mutants compared to the parental Δ*divJ* Δ*pleC* strain (Figure 3f). Furthermore, the activity of P_*pilA*_ and P_*hvyA*_ was much lower in Δ*divJ* Δ*pleC* Δ*cckN* and in Δ*divJ* Δ*pleC cckN*_*H47A*_ than in Δ*divJ* Δ*pleC* cells (Figure 3d). In addition, expression of *cckN in trans* from the xylose-inducible promoter P_*xylX*_ could fully complement loss of *cckN* in the Δ*divJ* Δ*pleC* Δ*cckN* mutant as this strain was again fully sensitive to both CbK and Cr30 infections (Supplementary Figure 3f). Together, these data suggest that *cckN* controls development by sustaining optimal CtrA activity, especially when PleC activity is compromised.

### Overexpression of *cckN* suppresses Δ*pleC* defects by enhancing CtrA activity

The results presented above suggested that *cckN* overexpression should lead to an increase of CtrA phosphorylation. To test this hypothesis, *cckN* was first slightly overexpressed in a Δ*pleC* mutant. As expected from the data presented above, *cckN* overexpression in Δ*pleC* cells was able to partially restore CbK and Cr30 phage sensitivity (Supplementary Figure 4a), motility (data not shown) as well as *hvyA* and *pilA* expression (Supplementary Figure 4b). Thus, CckN is capable of replacing PleC function under conditions where PleC is absent. However, *cckN* overexpression in a Δ*pleC* background did not restore attachment (Supplementary Figure 4c), likely because holdfast production required for irreversible attachment essentially relies on PleC kinase rather than phosphatase activity (15). Nevertheless, *cckN* inactivation in a *pleC*_*F778L*_ or Δ*pleC* Δ*cckN* background led to a slight decrease of attachment (Supplementary Figure 3b, Supplementary Figure 4c). These effects could be due to the decrease of CtrA~P, which results in reduced expression of *pilA* involved in primary attachment (29) and *hfa* genes involved in holdfast attachment (10). This is supported by the fact that overexpression of *cckN* in Δ*pleC* cells further decreased attachment (Supplementary Figure 4c).

Next, we measured *in vivo* phosphorylation levels of CtrA in wild-type cells strongly overexpressing *cckN*. The phosphorylation but not the abundance of CtrA was enhanced upon *cckN* overexpression (Figure 4a), in line with the data presented above (Figure 3b). In addition, overexpression of *cckN* generated filamentous cells arrested in G1 (Figure 4b) – likely due to hyperactive CtrA~P interfering with DNA replication initiation – which manifested in a drastically reduced number of viable cells (Figure 4c). This toxicity of *cckN* overexpression was not observed in cells with reduced CtrA activity (*cckATS1* and *ctrA401*) (1, 3) or abundance (*cpdR*_*D51A*_ and *cpdR*_*D51A*_ Δ*divK*) (22) (Figure 4c). Similarly, CckN variants that are predicted to lack kinase and phosphatase activities (CckN^K-P−^) – *cckN*_*H47N*_ and *cckN*_*T51R*_, the latter of which was designed based on the PleC^K-P−^ variant T561R (20) – were less toxic upon overexpression compared to wild-type CckN (Figure 4d). In contrast, overexpression 273 of CckN variants predicted to have lost only kinase activity (CckN^K-P+^) – *cckN*_*F212L*_, designed based on PleC_F778L_ (20), and *cckN*_*D195N*_, harbouring a mutation in the G1 box required for ATP binding – were still as toxic as the wild-type (Figure 4d). Thus, our data suggest that the major role of CckN *in vivo* is not to phosphorylate, but to dephosphorylate DivK and PleD, and thereby to stimulate CtrA activity.

**Figure 4:**
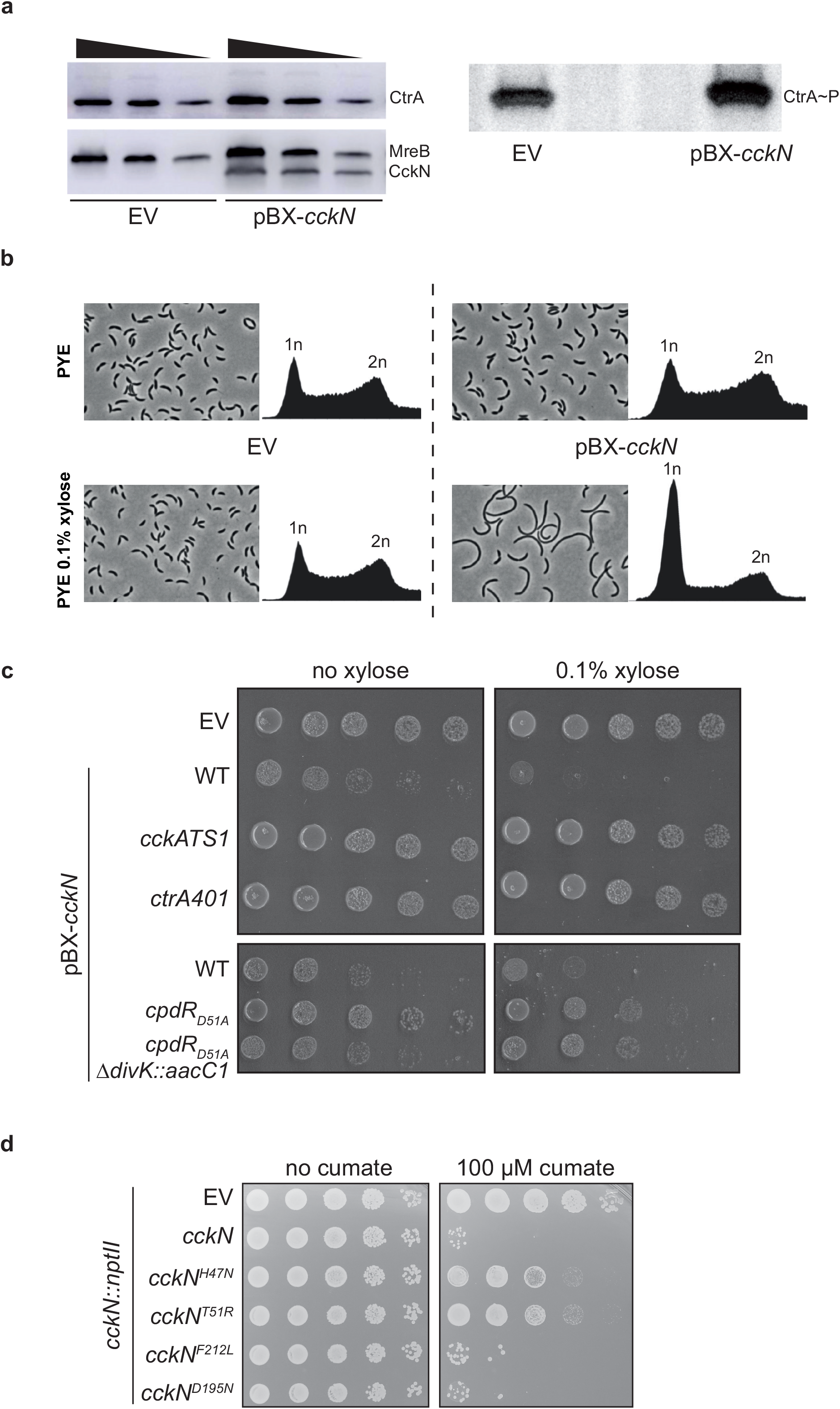
Overexpression of *cckN* leads to hyperactive CtrA~P and a subsequent toxic G1 block. (a) The protein (left) and phosphorylation (right) levels of CtrA were measured in wild-type (RH50) strain harbouring either a pBX or a pBX-*cckN* plasmid grown for 3 h in PYE (left) or M5G with 0.05 mM phosphate (right) supplemented with 0.1% xylose. Twofold serial dilutions of cell lysates were separated on SDS-PAGE to determine CtrA, MreB and CckN proteins levels by western blotting. The CtrA phosphorylation levels were determined *in vivo* as described in the Material & Methods. (b) Phase contrast imaging and FACS profiles of wild-type (RH50) cells harbouring either a pBX or a pBX-*cckN* plasmid grown for 3 h in PYE supplemented with 0.1% xylose. (c) Serial dilutions of the wild-type (RH50) strain harbouring the empty pBX vector (EV) and the wild-type (RH50), *cckATS1* (RH340), *ctrA401* (RH212), *cpdR*_*D51A*_ (RH347), *cpdR*_*D51A*_ Δ*divK∷aacC1* (RH1411) strains harbouring pBXMCS-2-*cckN* grown in PYE were spotted on PYE (left) or PYE supplemented with 0.1% xylose (right) and incubated for 2 days at 30° C. (d) Serial dilutions of the *cckN∷nptII* strain harbouring the empty pQF vector (EV) or pQF vector expressing *cckN* variants grown in PYE were spotted on PYE (left) or PYE supplemented with 100 μM cumate (right) and incubated for 2 days at 30° C.

### Transcription of *cckN* is stimulated by CtrA~P and (p)ppGpp

Since our ChIP-Seq data suggested that *cckN* might be a direct target of CtrA (Supplementary Table 1), we monitored the activity of P_*cckN*_*∷lacZ* fusion in mutant strains harbouring higher or lower CtrA activity. We found that the activity of P_*cckN*_ was decreased or increased in strains harbouring lower (Δ*pleC*, *ctrA401* and *cckATS1*) or higher (Δ*divJ* and *divK341*) CtrA activity, respectively (Supplementary Figure 5a). We also found that P_*cckN*_ activity was induced upon entry into stationary phase (Supplementary Figure 5b). In agreement with recent data showing that (p)ppGpp is required to sustain optimal activity of CtrA-dependent promoters during stationary phase (30), induction of P_*cckN*_ was not observed in (p)ppGpp^0^ (Δ*spoT*) cells (Supplementary Figure 5b). In addition, ectopic production of (p)ppGpp – from a functional truncated version of the *E. coli* (p)ppGpp synthetase RelA expressed from the xylose-inducible promoter (P_*xylX*_∷*relA*’) – increased P_*cckN*_ activity in exponentially growing cells (Supplementary Figure 5c). Together, these results suggest that induction of *cckN* expression in stationary phase by CtrA~P is (p)ppGpp-dependent.

### PleC and CckN are cleared from G1 cells in a ClpXP-dependent way

Based on the results presented above, PleC and CckN protect premature inactivation of CtrA during the G1 phase of the cell cycle, by maintaining the phosphorylation level of DivK low. At the G1-S transition, DivK~P levels raise to indirectly trigger CtrA dephosphorylation and ClpXP-dependent proteolysis. This implies that both CckN and PleC should be inactivated first. Thus, we tested whether CckN abundance fluctuates along the cell cycle by monitoring levels of CckN fused to a triple FLAG tag to either its N- or C-terminus. Interestingly, CckN-3FLAG was present only in G1/swarmer cells whereas 3FLAG-CckN remained roughly constant throughout the cell cycle (Figure 5a). The rapid disappearance of CckN-3FLAG (Supplementary Figure 6a-b) suggests the involvement of ATP-dependent proteolysis. To identify the protease responsible for CckN proteolysis, CckN-3FLAG abundance was quantified in a set of known *C. crescentus* protease and proteolytic adaptors mutant strains. As expected, the abundance of CtrA increased in Δ*clpX* and Δ*clpP* strains, as well as in strains lacking its known proteolytic adaptors CpdR, RcdA and PopA (9, 17, 31) (Figure 5b). Note that deletion of *clpX* and *clpP* genes were in a Δ*socAB* background that suppresses their essentiality (32). CckN-3FLAG levels increased in the Δ*clpX* and Δ*clpP* mutants, and this effect was independent of known ClpXP proteolytic adaptors (Figure 5b). In agreement with this result, we found that CckN-3FLAG levels properly fluctuated throughout the cell cycle in Δ*cpdR* cells, in contrast to CtrA whose degradation of which strictly depends on CpdR (Supplementary Figure 6d). As PleC abundance was also suggested to vary along the cell cycle (33) by disappearing together with CtrA at the G1-S transition (Supplementary Figure 6a-b), we also determined PleC abundance in the same protease and proteolytic adaptors mutants. Similar to CckN, PleC abundance increased in Δ*clpX* and in Δ*clpP* mutants but not in strains lacking CpdR, RcdA and PopA adaptors (Figure 5b). These results suggest that other proteolytic adaptors required for timely degradation of key cell cycle regulators remain to be uncovered.

**Figure 5:**
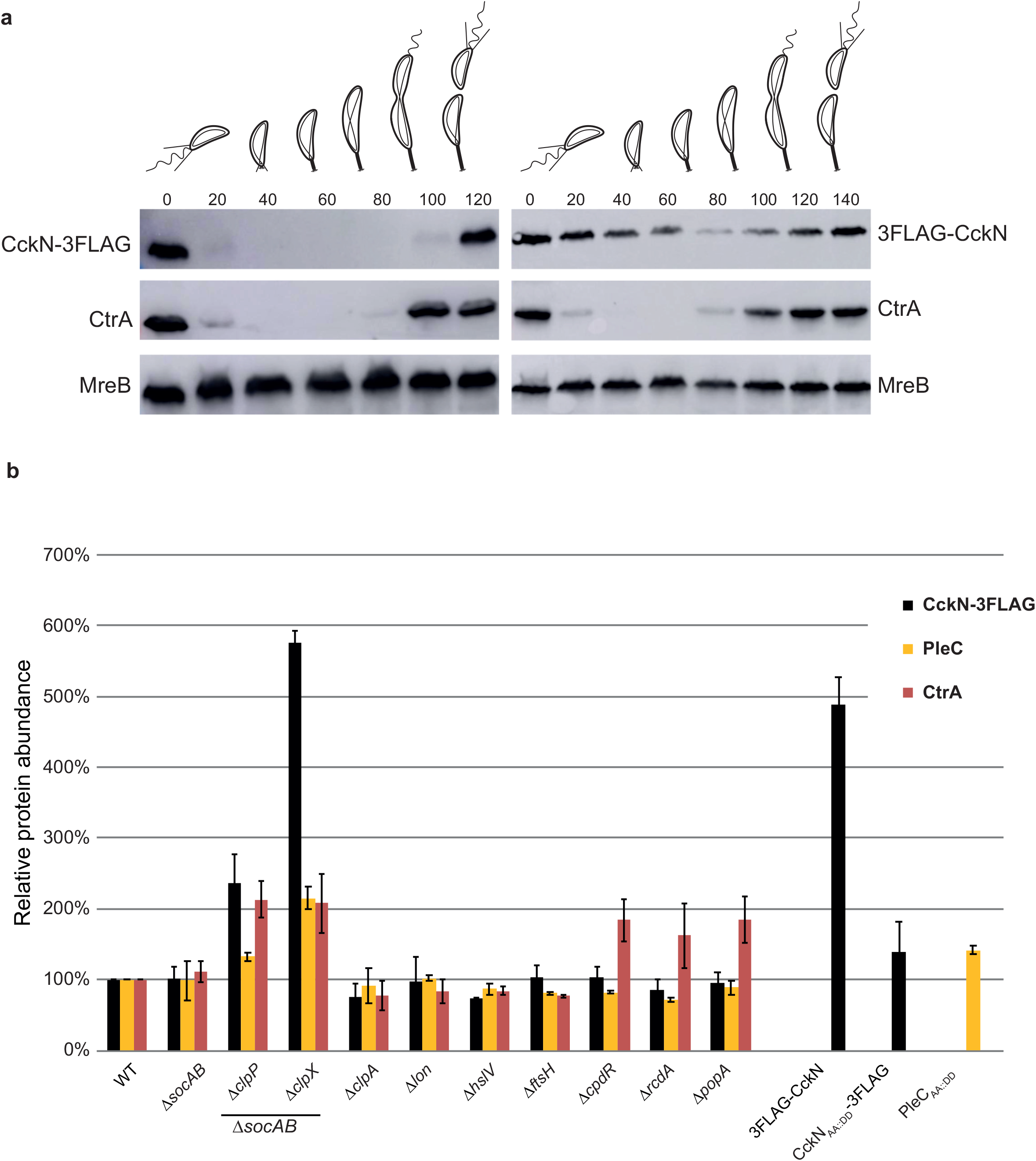
The ClpXP protease is responsible for the cell cycle oscillation of CckN and PleC. (a) Immunoblotting of protein samples extracted from synchronized *cckN-3FLAG* (RH1881) or *3FLAG-CckN* (RH1929) cells to follow CckN, CtrA and MreB abundance throughout the cell cycle. The time at which the samples were withdrawn after synchrony are indicated in minutes. (b) The relative abundance of CckN-3FLAG, PleC and CtrA was measured in wild-type (RH50 or RH1881), Δ*socAB* (RH1671 or RH737), Δ*socAB* Δ*clpP* (RH2279 or RH1063), Δ*socAB* Δ*clpX* (RH1674 or RH995), Δ*clpA* (RH864 or RH1093), Δ*lon* (RH2472 or RH1228), Δ*hslV* (RH864 or RH1247), Δ*ftsH* (RH865 or RH1229), Δ*cpdR* (RH339 or RH1133), Δ*rcdA* (RH323 or RH1149), Δ*popA* (RH315 or RH1151), *3FLAG-cckN* (RH1929), *cckN*_*AA∷DD*_*-3FLAG* (RH1576) and *pleC*_*AA∷DD*_ (RH2548) strains grown in complex medium (PYE) and normalized to the WT (100%). Error bars = SD, n = 3.

Previous studies demonstrated that substituting the two C-terminal hydrophobic residues of cell cycle regulators proteolyzed by ClpXP by two aspartate residues (*i.e.* CtrA_AA∷DD_, KidO_VA∷DD_, TacA_AG∷DD_, GdhZ_AA∷DD_ and ShkA_AG∷DD_) led to their stabilization (2, 34–37). Since both CckN and PleC displayed two alanine residues at their C-terminal extremity (Supplementary Figure 6c), we tested their potential involvement in the recognition of PleC and CckN by ClpX by creating *pleC*_*AA∷DD*_ and *cckN*_*AA∷DD*_*-3FLAG* mutants and monitoring protein abundance in asynchronous and synchronized populations. CckN_AA∷DD_-3FLAG was not protected from degradation along cell cycle (Figure 5b, Supplementary Figure 6e). In contrast, PleC_AA∷DD_ levels increased and did not oscillate anymore over the cell cycle (Figure 5b, Supplementary Figure 6f-g), suggesting that the C-terminal motif Ala-Ala is critical for PleC but nor for CckN to be recognized by ClpX. Given that a 3FLAG fusion at the N-terminal extremity of CckN led to protein stabilization whereas the same tag at the C-terminal end did not interfere with proteolysis (Figure 5a), a N-terminal motif instead of the two C-terminal hydrophobic residues is likely involved in the recognition of CckN by ClpX. Altogether, our data show that CckN and PleC are degraded in a ClpXP-dependent early in G1 phase, before the inactivation of the master regulator CtrA at the G1-S transition by dephosphorylation and proteolysis. However, in contrast to cell cycle regulators described so far to be proteolyzed by ClpXP, including CtrA itself, degradation of CckN and PleC likely requires unidentified proteolytic adaptors and involves – at least for CckN – an unsuspected N-terminal motif.

## Discussion

CckN was identified almost two decades ago in a yeast two-hybrid screen as a an interaction partner of the response regulator DivK (13). Here, we confirmed this interaction and also uncovered CckN as a second functional phosphatase for DivK and PleD (Figure 1). In contrast to the primary phosphatase PleC whose inactivation leads to pleiotropic effects, *cckN* loss-of-function mutants (Δ*cckN* or *cckN*_*H47A*_) did not display any detectable phenotype in an otherwise wild-type background. In contrast, deletion of *cckN* in a Δ*divJ* Δ*pleC* background led to a decrease of CtrA activity (Figure 3, Supplementary Figures 1 and 3, Supplementary Table 1) as well as developmental defects. In addition, a slight overexpression of *cckN* suppresses Δ*pleC* defects associated with its phosphatase, but not kinase, activity (Supplementary Figure 4) while a stronger overexpression leads to CtrA-dependent G1 block and toxicity (Figure 4). Thus, our data suggest that PleC phosphatase might be seconded by CckN upon specific conditions that still need to be uncovered.

Despite the redundancy of their phosphatase activity, CckN and PleC display different subcellular localization since PleC operates at the swarmer pole while CckN is diffused into the cytosol (Supplementary Figure 7). It is noteworthy that a CckN-GFP fusion expressed from the native *cckN* locus was almost undetectable, suggesting that the expression level of *cckN* is low. It is likely that the polar localization of PleC (11) has been selected for regulating its kinase rather its phosphatase activity. Indeed, PleC^K^ is allosterically activated at the differentiating pole by polar DivK~P (15), suggesting that the kinase activity of PleC is restricted to that pole. In contrast, PleC still displays phosphatase activity when not polarly localized, as shown in Δ*spmX* cells (28). In Δ*spmX*, the activity of the σ^54^-dependent transcriptional activator TacA is exacerbated (38). A comparative transposon (Tn) insertions coupled to deep sequencing (Tn-Seq) experiment done on Δ*spmX vs* wild-type cells supported that conclusion, since insertions into genes coding for positive regulators of TacA activity or expression were over-represented (38). This was the case for *shkA* coding for the hybrid kinase ShkA activating TacA by phosphorylation or *rpoN* encoding σ^54^ (38). Interestingly, Tn insertions also strongly accumulated in *pleC* and *cckN* in this experiment, likely because inactivation of their phosphatase activity down-regulates the CtrA-dependent P_*tacA*_ promoter, thereby limiting *tacA* expression. Alternatively, or in addition, a reduction in CtrA activity might also result in decreased expression of other genes affecting the ShkA-TacA pathway, such as *pleD* or other factors that could influence the phosphorylation state of the PleD/ShkA/TacA axis. Since TacA expression largely relies on the phosphatase activity of PleC (37), these Tn-Seq data support the redundancy between CckN and PleC. Notwithstanding, it has been shown that TacA phosphorylation by ShkA during the G1 phase requires c-di-GMP synthesized by PleD (37). Since TacA activity – monitored with a P_*spmX*_*∷lacZ* translational fusion – was not decreased in Δ*divJ* Δ*pleC* cells but strongly diminished in Δ*pleD* cells, it has been proposed that a third histidine kinase phosphorylates PleD in a Δ*divJ* Δ*pleC* background (37). In agreement with the fact that CckN does not display kinase activity (Supplementary Figure 1), we found that P_*spmX*_ activity was as high in Δ*divJ* Δ*pleC* Δ*cckN* cells as in wild-type cells (Supplementary Figure 8), thereby excluding CckN as the missing kinase for PleD. Thus, other phosphodonors for PleD, as well as for DivK as already suggested (11), remain to be uncovered. Indeed, the fact that *cckN* inactivation causes DivK-dependent phenotypes in a Δ*divJ* Δ*pleC* background demonstrates that, in the absence of DivJ and PleC, DivK is largely functional. The fact that CckN does seem to play a modulatory role on DivK/CtrA activity, rather than an essential, suggests the presence of yet other regulators affecting DivK.

PleC and DivJ are also able to modulate PleD phosphorylation (14, 15, 39). We show here that PleD is completely dephosphorylated *in vivo* upon *cckN* overexpression (Figure 2e), very likely as a result of two complementary effects. First, by decreasing phosphorylation levels of DivK, which was shown to allosterically affect DivJ and PleC kinase activity and thereby PleD phosphorylation (15). Second, by directly dephosphorylating PleD~P (Figure 2d). Concomitantly maintaining low PleD~P and DivK~P levels is important for coordinating cell cycle and developmental control. Indeed, strong activation of PleD in G1 cells would trigger premature abandon of motility without starting DNA replication. Interestingly, no autokinase activity was detected with CckN even in the presence DivK_D53N_, yet DivK_D53N_ was able to strongly induce kinase activity of DivJ and PleC (15) (Supplementary Figure 1).

Surprisingly, we observed phosphorylated CckN in the presence of DivJ and ATP (Supplementary Figure 1b). Thus, CckN and DivJ might interact with each other to form heterodimers in which CckN is trans-phosphorylated by DivJ. If true, this would even further protect DivK and PleD from premature activation during G1 or G1-S transition by draining phosphoryl-groups away from DivJ to CckN. CckN could act as a prototypical phosphatase, *i.e.* catalyzing hydrolysis of phospho-aspartate on receiver domain (REC-Asp~P) of DivK and PleD via coordination of a water molecule without being phosphorylated itself. Alternatively, CckN could serve as a sort of histidine phosphotransferase (Hpt) that accepts phosphoryl groups from REC-Asp~P in a back-transfer reaction, thereby leading to REC dephosphorylation while being phosphorylated itself. In fact, the phosphorylatable His residue required for autophosphorylation is dispensable for phosphatase activity in the prototypical bifunctional histidine kinase/phosphatase EnvZ, where His243 can be replaced by several amino acids still allowing significant dephosphorylation of its cognate substrate, OmpR (40). In contrast, His47 of CckN seems essential for its proposed function in dephosphorylating DivK and PleD, since replacement of His47 in CckN by Ala or Asn essentially phenocopies a *cckN* null allele in all assays tested. Of note, the idea that bifunctional histidine kinases/phosphatases with an intact CA domain can mainly function as Hpts to distribute phosphoryl groups between up- and downstream components is not unprecedented. It was recently proposed that LovK and PhyK involved in the general stress response in *C. crescentus* exert their main functions by acting as Hpts (41). PhyK and LovK share similarity in their DHp domains, but belong to different histidine kinase subfamilies, respectively HisKA2 and HWE, the former of which essentially has a CA domain identical to HisKA kinases with all known residues important for autophosphorylation conserved (41–43). PhyP from the Alphaproteobacterium *Sphingomonas melonis* Fr1 harbors a DHp domain similar to PhyK/LovK but fails to classify as either HisKA2 or HWE kinase due to its degenerate CA domain (44). PhyP was initially described as a phosphatase for PhyR, the response regulator universally controlling the general stress response in Alphaproteobacteria (44, 45). However, PhyP was recently shown to act as a HPt rather than a true phosphatase, shuttling phosphoryl group towards or away from PhyR (46). Thus, histidine kinases/phosphatases exist that employ back-transfer of phosphoryl groups as a means to dephosphorylate response regulator. Whether CckN (and PleC) belong to this group of enzymes employing such a mechanism remains to be studied in the future.

Our data strongly suggest that both CckN and PleC are proteolyzed in a ClpXP-dependent manner in early G1 cells (Figure 5, Supplementary Figures 6). Unexpectedly and in contrast to any ClpXP substrates described to date, it seems that none of the known proteolysis adaptors is required for CckN or PleC degradation. This is surprising knowing that CckN was found as a potential partner of RcdA in a pulldown assay (47). It is possible that CckN interacts with RcdA to limit its availability as a protease adaptor and anti-adaptor during early G1 phase, thereby avoiding premature RcdA functions in the hierarchical proteolytic events. Since both phosphatases disappear very early in G1 phase, prior to all other cell cycle regulators proteolytically degraded at the G1-S transition, a novel adaptor could be needed. Alternatively, proteolysis could be directly triggered *in vivo* by a direct interaction with ClpX without the help of an adaptor.

Since CckN is restricted to G1 cells in *C. crescentus*, its functions could be required in conditions leading to an extended G1 phase. In its natural oligotrophic environment, *C. crescentus* is expected to encounter extended periods of nutrient starvation during which the swarmer cells might not initiate DNA replication. Indeed, we know that limiting nitrogen or carbon leads to a (p)ppGpp-dependent G1 block in *C. crescentus* (48, 49) and that *cckN* expression is positively regulated by (p)ppGpp (Supplementary Figure 5b-c). In these stressful conditions, CckN might therefore be required to second PleC in maintaining DivK phosphorylation low and avoiding premature inactivation CtrA~P. We tested this hypothesis by comparing the viability of wild-type and Δ*cckN* cells maintained in stationary phase of growth. However, despite a (p)ppGpp-dependent induction of *cckN* expression in these conditions, Δ*cckN* cells remained as viable as the wild-type when maintained in stationary phase for days (Supplementary Figure 5d) or for weeks (data not shown). Likewise, we did not find any particular stressful conditions (nutrient starvation, oxidative stress, heavy metal exposure, temperatures, …) that decreased viability of Δ*cckN* more than wild-type cells, whether *pleC* was present or not, and whether the strains were incubated individually or mixed equally in the same culture (data not shown).

In alphaproteobacteria, DivK phosphorylation is known to be regulated by multiple paralogous histidine kinases, which based on similarity of their DHp domains belongs to the PleC/DivJ-like family, with numbers ranging from one in the obligate intracellular pathogen *Rickettsia prowazekii* to four in the facultative host-associated bacteria *Sinorhizobium meliloti* or *Agrobacterium tumefaciens* (50–56). Interestingly, the role played by these proteins on DivK phosphorylation varies between species. For instance, *B. abortus* PdhS is a *bona fide* bi-functional HK regulating positively and negatively the phosphorylation of DivK (57). In contrast, neither DivJ nor PleC was able to modulate DivK phosphorylation (Nayla Francis, personal communication) or localization (51). However, as both HK can physically interact with DivK, it has been hypothesized that DivJ and PleC could display kinase or phosphatase activity in specific conditions, such as in *Brucella*-containing vacuoles during cellular infection. In *S. meliloti*, DivK can be phosphorylated by DivJ and CbrA and dephosphorylated by PleC (54), whereas the function of PdhSA remains unknown. Thus, the multiple PleC/DivJ-like proteins that interact with DivK likely refer to the various environments encountered by these alphaproteobacteria depending on their lifestyles. The challenge now will be to identify the specific stimuli regulating the activity of these DivK-associated HK.

## Material & Methods

### Bacterial strains and growth conditions

*Escherichia coli* Top10 was used for cloning purpose, and grown aerobically in Luria-Bertani (LB) broth (Sigma). Electrocompetent or thermocompetent cells were used for transformation of *E. coli.* All *Caulobacter crescentus* strains used in this study are derived from the synchronizable wild-type strain NA1000, and were grown at 30 °C in Peptone Yeast Extract (PYE) or synthetic media supplemented with glucose (M2G or M5G) as already described in (49). Genes expressed from the inducible *xylX* promoter (P_*xylX*_) was induced with 0.1% to 0.2% xylose in *xylX*^*+*^ background. Motility was assayed on PYE plates containing 0.3% agar. Generalized transduction was performed with phage Cr30 according to the procedure described in (58).

For *E. coli*, antibiotics were used at the following concentrations (μg/ml; in liquid/solid medium): ampicillin (100/100), kanamycin (30/50), oxytetracycline (12.5/12.5), spectinomycin (100/100), streptomycin (50/100) or chloramphenicol (20/30) For *C. crescentus*, media were supplemented with kanamycin (5/20), oxytetracycline (2.5/2.5), spectinomycin (25/50), streptomycin (5/5), hygromycin (100/100), nalidixic acid (20), chloramphenicol (1/2) or gentamycin (0.5/5) when appropriate. Cumate (4-isopropylbenzoic acid) was dissolved in 100% ethanol to result in a 1000X 100 mM stock solution and used at a final concentration of 100 uM in plates. For “no cumate” controls, an equal volume of ethanol was added to plates.

### Bacteriophage sensitivity assays

The sensitivity to CbK (LHR1) and CR30 (LHR2) bacteriophages were performed as follows. First, 200 μl overnight culture of *Caulobacter* cells grown in PYE were mixed to 4-5 ml of prewarmed PYE Top Agar (0.45% agar) medium and poured on a PYE plain agar plate. Then, CbK and Cr30 bacteriophage lysates were serially diluted spotted (5 μl drops) on the Top Agar once solidified and incubated overnight at 30 °C.

### Construction of plasmids and strains

Detailed descriptions of bacterial strains are included in the Supplementary Information. Strains, plasmids and oligonucleotides used for plasmids and strains construction are listed in Tables S1-S3. *E. coli* S17-1 and *E. coli* MT607 helper strains were used for transferring plasmids to *C. crescentus* by respectively bi- and tri-parental matings. In-frame deletions were created by using the pNPTS138-derivative plasmids as follows. Integration of the plasmids in the *C. crescentus* genome after single homologous recombination were selected on PYE plates supplemented with kanamycin. Three independent recombinant clones were inoculated in PYE medium without kanamycin and incubated overnight at 30 °C. Then, dilutions were spread on PYE plates supplemented with 3% sucrose and incubated at 30 °C. Single colonies were picked and transferred onto PYE plates with and without kanamycin. Finally, to discriminate between mutated and wild-type loci, kanamycin-sensitive clones were tested by PCR on colony using locus-specific oligonucleotides.

### Attachment assays

Overnight cultures were diluted in a 96-well microplate and incubated for 18 h to 24 h at 30 °C before measuring absorbance at the optical density of 660 nm (OD_660_). Unattached cells were discarded and the microplate was washed 3 times with dH_2_O. Then, 0.1% crystal Violet (CV) was added for 15 min under agitation before washing the wells with dH_2_O. Finally, CV was dissolved in a 20% acetic acid solution for 15 min under agitation and absorbance at 595 nm (OD_595_) was taken. To normalize attachment to growth, the ratio between OD_595_ and OD_660_ was used.

### Spotting assays

For experiments shown in Figure 4c-d, 10-fold serial dilutions (in PYE) were prepared in 96-well plates from 5 ml cultures in standard glass tubes grown overnight at 30 °C in PYE Kan (Figure 4c) or Tet (Figure 4d) and dilution series were spotted on PYE Kan and PYE Kan 0.1% xylose (Figure 4c) or replica-spotted on PYE Tet and PYE Tet Q100 plates using an in-house-made 8-by-6 (48-well) metal pin replicator (Figure 4d). Plates were incubated at 30 °C for two days and pictures were taken.

### Synchronization of cells

Synchronization of cells was performed as already described in (36). Briefly, cells were grown in 200 ml of PYE to OD_660_ of 0.6, harvested by centrifugation for 20 min 556 at 5,000 × g, 4 °C; resuspended in 60 ml of ice-cold Phosphate (PO_4_^3−^) buffer and combined with 30 ml of Ludox LS Colloidal Silica (30%) (Sigma-Aldrich). Cells resuspended in Ludox was centrifuged for 40 min at 9,000 × g, 4 °C. Swarmer cells, corresponding to the bottom band, were isolated, washed twice in ice-cold PO_4_^3−^ buffer, and finally resuspended in prewarmed PYE media for growth at 30 °C. Samples were collected every 15 min for western blot, microscopy and FACS analyses.

### Proteins purification

In order to immunize rabbits for production of polyclonal antibodies and or to perform *in vitro* phosphorylation assays, DivJ^Sm^, CckN, DivK and His6-PleC_505-842_ was purified as follows. A BL21 (DE3) strain harbouring plasmid pML31-TRX-His6-*divJ*^*HK*^_*Sm*_ (54), pET-28a-*cckN*, pET-28a-*divK* or pET-28a-*pleC*_*505-842*_ was grown in LB medium supplemented with kanamycin until OD_600_ of 0.7. IPTG was added at a final concentration of 1 mM and the culture was incubated at 37 °C for 4 h. Then, cells were harvested by centrifugation for 20 min at 5,000 × g, 4 °C. The pellet was resuspended in 20 ml BD buffer (20 mM Tris-HCl pH 8.0, 500 mM NaCl, 10% glycerol, 10 mM MgCl_2_, 12.5 mM Imidazole) supplemented with complete EDTA-free protease cocktail inhibitor (Roche), 400 mg lysozyme (Sigma) and 10 mg DNaseI (Roche) and incubated for 30 min on ice. Cells were then lysed by sonication and the lysate by centrifugation 12,000 rpm for 30 min at 4°C was loaded on a Ni-NTA column and incubated 1 h at 4 °C with end-over-end shaking. The column was then washed with 5 ml BD buffer, 3 ml Wash1 buffer (BD buffer with 25 mM imidazole), 3 ml Wash2 buffer (BD buffer with 50 mM imidazole), 3 ml Wash3 buffer (BD buffer with 75 mM imidazole). Proteins bound to the column were eluted with 3 ml Elution buffer (BD buffer with 100 mM imidazole) and aliquoted in 300 μl fractions. All the fractions containing the protein of interest (checked by Coomassie blue staining) were pooled and dialyzed in Dialysis buffer (50 mM Tris pH 7.4, 12.5 mM MgCl_2_).

For experiments shown in Figure 2d and Supplementary Figure 2, proteins were expressed in *E. coli* BL21 (DE3) harbouring plasmid pETHisMBP-*cckN*, pETHisMBP-*divJ* or pETHisMBP-*pleC*, or in *E. coli* BL21 (DE3) pLys harboring plasmidspCC2 or pRP112. Strains were grown overnight in 5 ml LB-Miller at 37 °C with appropriate antibiotics, diluted 100-fold in 500 ml LB-Miller with appropriate antibiotics and grown at 37 °C until the cultures reached an OD_600_ of 0.5–0.8. Cultures were then shifted to 23 °C and incubated for another hour, after which protein expression was induced by addition of 0.2 mM IPTG. After incubation overnight, cells were harvested by centrifugation (5000 × g, 20 min, 4 °C), washed once with 20 ml of 1X PBS, flash-frozen in liquid N2 and stored at −80 °C until purification. For purification, the pellet was resuspended in 8 ml of buffer A (2X PBS containing 500 mM NaCl, 10 mM imidazole and 2 mM β-mercaptoethanol) supplemented with DNaseI (AppliChem) and Complete Protease inhibitor (Roche). After one passage through a French press cell, the suspension was centrifuged (10,000 × g, 30 min, 4 °C) and the supernatant was mixed with 800 μl of Ni-NTA slurry, prewashed with buffer A, and incubated for 1-2 h on a rotary wheel at 4 °C. Ni-NTA agarose was loaded on a polypropylene column and washed with at least 50 ml of buffer A, after which the protein was eluted with 2.5 ml of buffer A containing 500 mM imidazole. The eluate was immediately loaded on a PD-10 column pre-equilibrated with kinase buffer (10 mM HEPES-KOH pH 8.0, 50 mM KCl, 10% glycerol, 0.1 mM EDTA, 5 mM MgCl_2_, 5 mM β-mercaptoethanol). The protein was then eluted with 3.5 ml of kinase buffer and stored at 4 °C until further use. All experiments were performed within one week after purification.

### Immunoblot analysis

Proteins crude extracts were prepared by harvesting cells from exponential growth phase, resuspending the pellet in SDS-PAGE loading buffer and lysing cells by incubating them for 10 min at 95 °C. Proteins were then subjected to electrophoresis in a 12% SDS-polyacrylamide gel, transferred onto a nitrocellulose membrane then blocked overnight in 5% (wt/vol) nonfat dry milk in phosphate buffer saline (PBS) with 0.05% Tween 20. Membrane was immunoblotted for ≥ 3 h with primary antibodies : anti-M2 (1:5,000) (Sigma), anti-CtrA (1:5,000), anti-MreB (1:5,000), anti-PleC (1:5,000), anti-CckN (1:2,000), anti-DivK (1:2,000) then followed by immunoblotting for ≤ 1 h with secondary antibodies: 1:5,000 anti-mouse (for anti-FLAG) or 1:5000 anti-rabbit (for all the others) linked to peroxidase (Dako Agilent), and vizualized thanks to Clarity™ Western ECL substrate chemiluminescence reagent (BioRad) and Amersham Imager 600 (GE Healthcare).

### FACS analysis

Bacterial cells were incubated at 30 °C until they reached mid-log phase, and 100 μL cells was added to 900 μl ice-cold 70% (vol/vol) ethanol solution and stored at −20 °C for 4 h or until further use. For rifampicin treatment, the mid-log phase cells were grown in the presence of 20 μg/mL rifampicin at 30 °C for 3 h. One milliliter of these cells was fixed as mentioned above. For staining and analysis, 2 ml fixed cells were pelletized and washed once with 1 ml staining buffer (10 mM Tris·HCl pH 7.2, 1 mM EDTA, 50 mM sodium citrate + 0.01% TritonX-100). Then, cells were resuspended in 1 ml staining buffer containing 0.1 mg/ml RNaseA (Roche Life Sciences) and incubated at room temperature (RT) for 30 min. The cells were then harvested by centrifugation at 6,000 × g for 2 min, and the pellets were resuspended in 1 ml staining buffer supplemented with 0.5 μM Sytox Green Nucleic Acid Stain (Molecular Probes, Life Technologies), followed by incubation in the dark for 5 min. These cells were then analyzed immediately in flow cytometer (FACS Calibur, BD Biosciences) at laser excitation of 488 nm. At least 1 × 10^4^ cells were recorded in triplicate for each experiment.

### Microscopy

All strains were imaged during exponential phase of growth (OD_660_ between 0.1 and 0.4) after immobilization on 1.5% PYE agarose pads. Microscopy was performed using Axioskop microscope (Zeiss), Orca-Flash 4.0 camera (Hamamatsu) and Zen 2.3 software (Zeiss). Images were processed using ImageJ.

### *In vivo* ^32^P labeling

A single colony of cells picked from a PYE agar plate was washed with M5G medium lacking phosphate and was grown overnight in M5G with 0.05 mM phosphate to OD_660_ of 0.3. One milliliter of culture was then labeled for 4 min at 30 °C using 30 μCi γ-[^32^P]ATP. Following lysis, proteins were immunoprecipitated with 3 μl of polyclonal anti-sera (anti-CtrA or anti-PleD). The precipitates proteins were resolved by SDS-PAGE and [^32^P]-labeled proteins was visualized using a Super Resolution screen (Perkin Elmer and quantified using a Cyclone Plus Storage Phosphor System (Perkin Elmer). The signal was normalized to the relative cellular content as determined by immunoblotting of whole cell lysates probed with antibodies.

### *In vitro* phosphorylation assays

For experiments shown in Figure 1c, autophosphorylation was performed on 5 μM of DivJ_Sm_ kinase in 50 mM Tris-HCl pH 7.4 supplemented with 2 mM DTT, 5 mM MgCl_2_, 500 μM ATP and 2 μCi γ-[^32^P]ATP at 30°C for 40 min. Then, 5 μM of DivK were added to the mix supplemented with 5 mM MgCl_2_ and phosphotransfer was performed at RT for 15 min. To remove excess of ATP, sample was washed twice in amicon column with a cut-off of 10 KDa by adding 450 μl of 50 mM Tris-HCl pH 7.4 and centrifuging 10 min at 12,000 rpm. CckN was then added at a final concentration of 5 μM and the mix was incubated 1, 2 and 5 min at RT. Reaction was stopped by adding SDS-PAGE loading buffer and samples were resolved by SDS-PAGE, the dried gel was visualized using a Super Resolution screen (Perkin Elmer) and quantified using a Cyclone Plus Storage Phosphor System (Perkin Elmer).

For experiments shown in Figure 2d and Supplementary Figure 2, all reactions were performed in kinase buffer supplemented with 433 μM ATP and 5-20 μCi γ-[^32^P]ATP (3000 Ci/mmol, Hartmann Analytic) at room temperature. For experiments shown in Supplementary Figure 2a, reactions containing His6-MBP-PleC (5 μM), His6-MBP-DivJ (5 μM) or His6-MBP-CckN (5 μM) without or with DivK_D53N_ (10 μM) were prepared and incubated for 20 min at RT before autophosphorylation was started by addition of ATP. Aliquots were withdrawn from the reactions as indicated in the figure and stopped by addition of 5X SDS sample buffer and stored on ice. For experiments shown in Supplementary Figure 2b, reactions containing His6-MBP-DivJ (9 μM), His6-MBP-CckN (11 μM), PleD (28 μM) and/or DivK_D53N_ (28 μM) were prepared and incubated for 30 min at RT before autophosphorylation was started. Reactions were stopped after 60 min by addition of 5X SDS sample buffer and stored on ice. For reactions shown in Figure 2d, His-MBP-DivJ (10 μM) was autophosphorylated for 60 min in a reaction volume of 150 μl, then 20 μL of PleD (8 μM final) were added, the reaction was split in 40 μl aliquots and, after allowing PleD phosphorylation for 2 min, 10 μl of His6-MBP-CckN (4 μM final) or kinase buffer (control) were added. Aliquots were withdrawn from the reactions as indicated in the figure and stopped by addition of 5X SDS sample buffer and stored on ice. Reactions were run on precast Mini-Protean TGX (Biorad) gels, wet gels were exposed to a phosphor screen, which was subsequently scanned using a Typhoon FLA 7000 imaging system (GE Healthcare).

### β-galactosidase assays

Overnight grown *Caulobacter* cells harboring lacZ reporter plasmids were incubated at 30 °C until OD_660_ of 0.3 to 0.5. Fifty microliters aliquots of the cells were treated with few drops of chloroform. To this, 750 μl of Z buffer (60 mM Na_2_HPO_4_, 40 mM NaH_2_PO_4_, 10 mM KCl, 1 mM MgSO_4_, pH 7.0) was added, followed by 200 μl of 4 mg/ml O-nitrophenyl-*β*-D-galactopyranoside (ONPG). Then, reaction was incubated at 30 °C until yellow color was developed, stopped by addition of 500 μl of 1 M Na_2_CO_3_ and OD_420_ of the supernatant was measured. The activity of the β-galactosidase expressed in miller units (MU) was calculated using the following equation: MU = (OD_420_ × 1,000) / [OD_660_ × t × v] where “t” is the time of the reaction (min), and “v” is the volume of cultures used in the assays (ml). Experimental values were the average of three independent experiments.

### Bacterial two-hybrid (BTH) assays

BTH assays were performed as described previously in (49). Briefly, 2 μl of MG1655 *cyaA∷frt* (RH785) strains expressing T18 and T25 fusions were spotted on MacConkey Agar Base plates supplemented with ampicillin, kanamycin, maltose (1%) and IPTG (1 mM), and incubated for 3-4 days at 30 °C. All proteins were fused to T25 at their N-terminal extremity (pKT25) or to the T18 at their N- (pUT18C) or C-terminal (pUT18) extremity. The β-galactosidase assays were performed on 50 μl *E. coli* BTH strains cultivated overnight at 30° C in LB medium supplemented with kanamycin, ampicillin and IPTG (1 mM) as described in (59).

### Co-immunoprecipitation (Co-IP) assays

*C. crescentus* cells were grown in 200 ml of PYE (supplemented with 0.1% xylose if required) to OD_660_ of 0.7 to 0.9), harvested by centrifugation for 20 min at 5,000 × g, 4°C. The pellets were washed once with PBS and resuspended in 10 ml PBS containing 2 mM DTSP (Dithiobis(succinimidyl propionate)) for crosslinking. After 30 min at 30°C, cross-linking was quenched by the addition of Tris-HCl to a final concentration of 0.150 M for 30 minutes. Cells were then washed twice with PBS and once with 20 ml Co-IP buffer (20 mM HEPES, 150 mM NaCl, 20% glycerol, 80 mM EDTA). The pellets were resuspended in lysis buffer [1x CelLytic B (Sigma), 10 mM MgCl_2_, 67.5U Ready-Lyse lysozyme (Epicentre), 10U/mL DNase I, 2% NP-40 Surfact-Amps Detergent (Thermo Scientific), ½ tablet Complete EDTA-free anti-proteases (Roche)] and incubated under soft agitation at RT for 30 min. Cells were then lysed by sonication, lysates were cleared by centrifugation and incubated 2 h at 4 °C with MagStrep type3 XT beads (Iba). The beads were washed 6 times with W Buffer (Iba) and precipitated proteins were released by 15 min incubation in BX Buffer (Iba). SDS loading buffer was added to samples and boiled for 10 min. Equal volumes of coimmunoprecipitates, and cell lysates from equal numbers of cells, were analyzed by SDS-PAGE and Western blotting. Membranes were probed with anti-DivK and anti-CckN primary antibodies.

### Chromatin immunoprecipitation followed by deep sequencing (ChIP-Seq) assays

ChIP-Seq assays were performed as described previously in (60). Briefly, 80 ml of mid-log phase cells (OD_600_ of 0.6) were cross-linked in 1% formaldehyde and 10 mM sodium phosphate (pH 7.6) at room temperature for 10 min and 30 min on ice thereafter. Crosslinking was stopped by addition of 125 mM glycine and incubated for 5 min on ice. Cells were washed thrice in PBS, resuspended in 450 μl in TES buffer (10 mM Tris-HCl pH 7.5, 1 mM EDTA, 100 mM NaCl) and lysed with 2 μl of Ready-lyse lysozyme solution for 5 min at RT. Protease inhibitors (Roche) was added for 10 min. Then, 550 μl of ChIP buffer (1.1% triton X-100, 1.2 mM EDTA, 16.7 mM Tris-HCl pH 8.1, 167 mM NaCl, plus protease inhibitors) was added to the lysate and incubated at 37 °C for 10 min before sonication (2 × 8 bursts of 30 sec on ice using a Diagenode Bioruptor) to shear DNA fragments to a length of 300 to 500 bp. Lysate was cleared by centrifugation for 10 min at 12,500 rpm at 4 °C and protein content was evaluated by measuring OD_280_. Then, 7.5 mg of proteins were diluted in ChIP buffer supplemented with 0.01% SDS and precleared 1 h at 4 °C with 50 μl of protein A agarose beads (BioRad) and 100 μg BSA. Two μl of polyclonal anti-CtrA antibodies were added to the supernatant before overnight incubation at 4 °C under gentle agitation. Eighty μl of BSA pre-saturated protein A agarose beads were added to the solution for 2 h at 4 °C with rotation, washed once with low salt buffer (0.1% SDS, 1% Triton X-100, 2 mM EDTA, 20 mM Tris-HCl pH 8.1, 150 mM NaCl), once with high salt buffer (0.1% SDS, 1% Triton X-100, 2 mM EDTA, 20 mM Tris-HCl pH 8.1, 500 mM NaCl), once with LiCl buffer (0.25 M LiCl, 1% NP-40, 1% deoxycholate, 1 mM EDTA, 10 mM Tris-HCl pH 8.1), once with TE buffer (10 mM Tris-HCl pH 8.1, 1 mM EDTA) at 4 °C and a second wash with TE buffer at RT. The DNA-protein complexes were eluted twice in 250 μl freshly prepared elution buffer (0.1 M NaHCO_3_, 1% SDS). NaCl was added at a concentration of 300 mM to the combined eluates (500 μl) before overnight incubation at 65 °C to reverse the crosslink. The samples were treated with 20 μg of proteinase K in 40 mM EDTA and 40 mM Tris-HCl (pH 6.5) for 2 h at 45 °C. DNA was extracted using Nucleospin PCR clean-up kit (Macherey-Nagel) and resuspended in 50 μl elution buffer (5 mM Tris-HCl pH 8.5). DNA sequencing was performed using Illumina HiSeq4000 (Genomicscore KULeuven and Bio.be Gosselies).

### NGS data analysis

Around 2 × 10^7^ single-end sequence reads (1 × 50) were first mapped against the genome of *C. crescentus* NA1000 (NC_011916.1) and converted to SAM using BWA (61) and SAM (62) tools respectively from the sourceforge server (https://sourceforge.net/). MACS2 (63) algorithm was used to model the length of DNA fragment as well as the shift size. Next, the number of reads overlapping each genomic position was computed using custom Python scripts and the previously modeled DNA fragment and shift sizes. A peak was defined as the genomic region where each position has more reads than the 97^th^ percentile. The candidate peaks were annotated using custom Python scripts. In the purpose to compare strains, the total number of reads was normalized by the ratio of the number of reads between the two strains.

## Acknowledgements

We are grateful to Patrick Viollier and Peter Chien for providing strains and plasmids; Emanuele Biondi for anti-CtrA antibodies; Fanny Zola and Aurélie Mayard for help with proteins purification. We thank the members of the BCcD team for critical reading of the manuscript and helpful discussions. The work conducted in the laboratory of U.J. was supported by the Swiss National Science Foundation (grants 31003A_166503 and 310030B_185372). This work was supported by a Research Credit (CDR J.0169.16) from the Fonds de la Recherche Scientifique – FNRS (F.R.S. – FNRS) to R.H. and R.H. is a Research Associate of F.R.S. – FNRS.

## Author Contributions

J.C., A.K., U.J. and R.H. conceived and designed the experiments. J.C. performed all the experiments except otherwise stated. A.K. performed *in vitro* phosphorylation assays shown in Figure 2d and Supplementary Figure 1 as well as the growth assays upon overexpression of *cckN* variants shown in Figure 4d. K.P. designed the bioinformatic tool to analyse the ChIP-Seq data. T.B. characterized the ClpXP-dependent degradation of PleC (Figure 5b and Supplementary Figure 6f-g). J.C., A.K., U.J. and R.H. analyzed the data. J.C., A.K. and R.H wrote the paper.

